# De novo designed inhibitor confers protection against lethal toxic shock

**DOI:** 10.1101/2024.08.23.608890

**Authors:** Robert J. Ragotte, Huazhu Liang, Jacob M. Berman, Matthias Glögl, Daniel Schramek, Roman A. Melnyk, David Baker

## Abstract

*Paeniclostridium sordellii* causes a toxic shock syndrome with a mortality rate of nearly 70%, primarily affecting postpartum and post-abortive women. This disease is driven by the production of the *P. sordellii* lethal toxin, TcsL, for which there are currently no effective treatments. We used a protein diffusion model, RFdiffusion, to design high affinity TcsL inhibitors. From a very small set of 48 starting designs and 48 additional sequence optimized designs, we developed a potent inhibitor with <100 pM affinity that protects mice prophylactically and therapeutically (post exposure) from lung edema and death in a stringent lethal challenge model. This inhibitor, which can be lyophilized without any loss of activity, is a promising therapeutic candidate for this rare but deadly disease, and our results highlight the ability of deep learning-based protein design to rapidly generate biologics with potential clinical utility.

## Main Text

*Paeniclostridium sordellii* lethal toxin, TcsL, is produced during *P. sordellii* infection and causes a highly lethal disease in humans and livestock. In humans, this primarily affects postpartum and post-abortive women, the latter associated with the use of misoprostol/mifepristone^1^. Other at-risk groups include intravenous drug use and those recovering from surgery or other traumatic injury^2^. The high (∼70%) lethality highlights a lack of effective therapeutic intervention for this rare disease^1,3–5^. Current treatments focus on clearing the infection with surgical removal of the affected tissue and the administration of broad spectrum antibiotics to eliminate the pathogen^3,6^, rather than neutralizing the toxin. The importance of developing new antitoxin therapies to improve clinical outcomes for *P. sordellii* toxic shock syndrome has long been recognized^3,5–7^, but little progress has been made to develop more targeted therapies.

TcsL uses a single known receptor binding site to engage with SEMA6A and SEMA6B on the host cell surface (Figure 1A)^8,9^. We reasoned that a drug that could bind to the toxin at that site and induce immediate cessation of TcsL activity could protect against disease progression and death. Given a protein target, the generative AI denoising diffusion network, RFdiffusion^10^, can build proteins from scratch that have high target shape complementarity, and following sequence design with ProteinMPNN^11^, have high binding affinity. Such small proteins can have advantages over monoclonal antibodies in ease of production and thermostability, while retaining high affinity and specificity for their targets^12–15^. We set out to design small proteins that could block the interaction of TcsL with its host cell receptors to protect against TcsL toxic shock.

**Figure 1.**
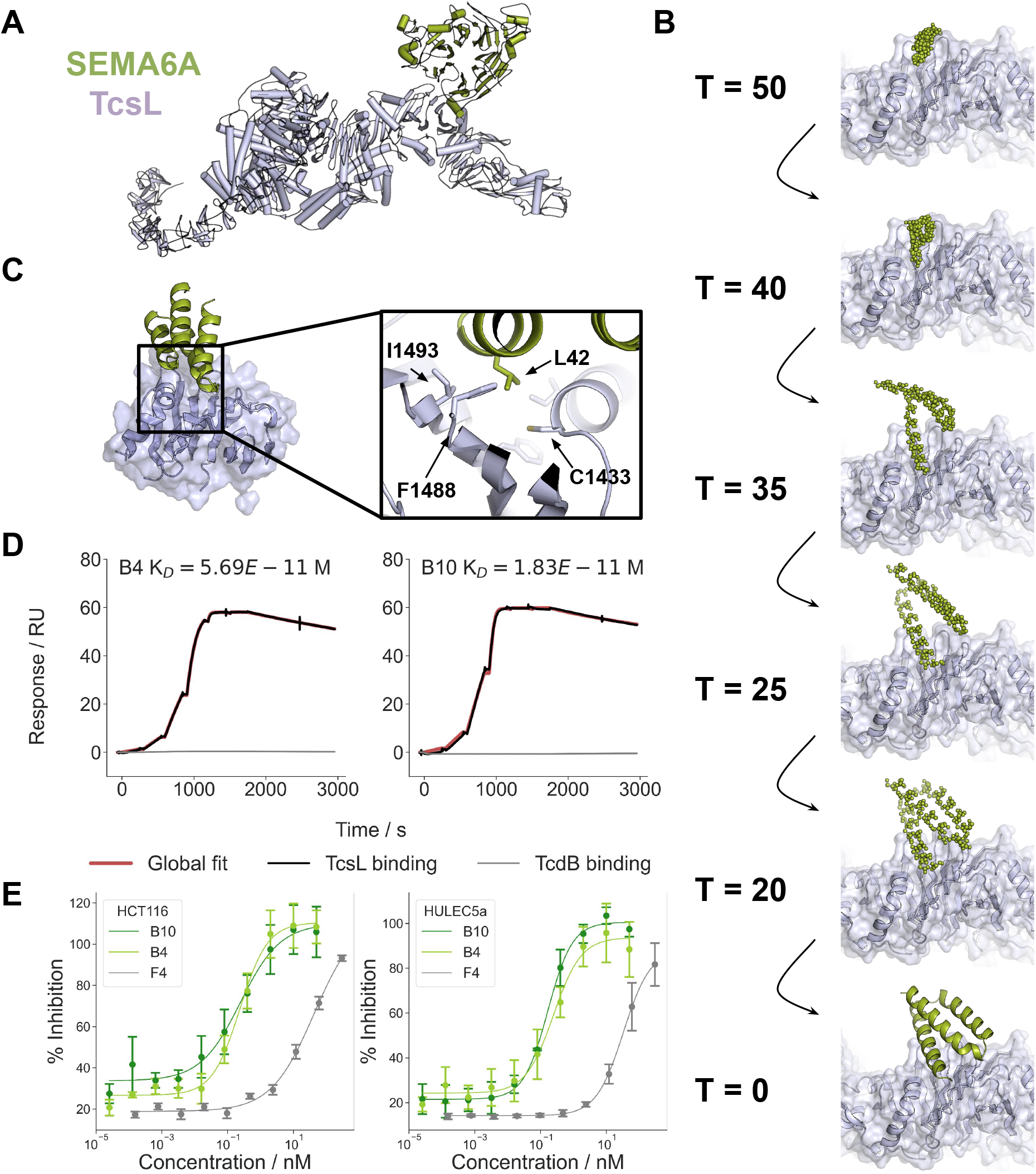
Design and optimization of TcsL blocking minibinders. **A**. Composite cartoon model of full length TcsL (8JB5) bound to SEMA6A (6WTS). **B**. An example of an RFdiffusion denoising trajectory starting from random noise placed above the target site at T=50 to the fully denoised design at T=0. **C**. Design model of B4 binding to the RBD of TcsL. The high degree of shape complementarity between the designed binder and the target is highlighted in the upper inset box. The lower inset box shows the hydrophobic pocket that accounts for the bulk of the interface. **D**. Single cycle kinetic analysis with SPR showing binding of two of the optimized designs to TcsL (black) and TcdB (gray) with toxin RBD captured and a 6-step 4-fold dilution series of the minibinder starting at 100 nM. **E**. Neutralization assays of parental (F4) and optimized (B4 and B10) designs using TcsL on HCT116 cells (left) and HULEC5a cells (right). Points indicate the mean and error bars are the standard error of the mean for n =3 (HULEC5a) and n = 4 (HCT116) independent replicates. (see main text for toxin concentrations).

We designed 55-65 residue miniprotein binders to the SEMA6A/B interface of TcsL using the denoising diffusion network RFdiffusion beginning from randomly generated residue coordinates and then iteratively denoising over 50 time steps (a representative denoising trajectory is shown in Figure 1B)^10^. 10,000 diffusion trajectories were targeted to hydrophobic hotspot residues at the SEMA6A interface (Figure 1C). Sequences that could fold into the target structure and were compatible with target site binding were designed using ProteinMPNN^11^ and the resulting designs filtered using AlphaFold2 initial guess^16^.

48 designs were produced in *E. coli* and screened for TcsL binding. The highest affinity, F4, bound with a K_D_ of 4 nM (Figure S1). F4 was further optimized using ProteinMPNN (see Methods)^16^ and 48 additional designs were experimentally tested (Figure S2). Two – B4 and B10 – were found to have very high affinities of 57 and 18 pM (Figure 1D), though 11 others had subnanomolar binding affinity to the RBD. These affinities are particularly remarkable as only 96 designs were tested in total, highlighting the ability of RFdiffusion to generate shape matching high affinity binders with relatively little experimental screening. For both B4 and B10, as well as the F4 parent, the bulk of the interface is centered around a hydrophobic pocket on the target site composed of C1433, I1434, I1437 and F1488 into which L42 on the binder extends (Figure 1C). The ProteinMPNN optimized designs contain P>K and S>K substitutions that likely form salt bridges with neighboring glutamate residues on TcsL.

In a viability assay using HCT116 cells and 50 pM of TcsL (the EC99 of the toxin on this cell line), the IC50s for B4 and B10 were 238 (95% CI 122 - 476) and 210 (95% CI 51.6 - 8171) pM, respectively (Figure 1E). Given that the primary target of TcsL during infection is lung vascular endothelial cells^17^, we tested neutralization using the HULEC5a cell line which has elevated SEMA6A expression. Using 0.5 pM TcsL (the toxin EC90 on this cell line), B4 and B10 remained highly protective with IC50s of 248 (95% CI 88 - 706) and 157 (95% CI 85 - 292) pM (Figure 1E). The designs bound specifically to TcsL, with no detectable binding to TcdB, despite 76% sequence identity between the two toxins (Figure 1D).

Given the success of these designs at inhibiting TcsL *in vitro*, we sought to evaluate these as therapeutic candidates in a mouse model of *P. sordellii* toxic shock. In an initial study, mice were administered TcsL alone or TcsL that was preincubated with 1000x molar excess of design B4. However, in this treatment regime, the molecules failed to protect the mice against disease (Figure S3).

We hypothesized that the reason for this was rapid glomerular filtration of the minibinder that was not bound to the toxin, given the small size of these biologics (∼9 kDa). To resolve this issue, we fused minibinder B4 to the albumin binding domain M79^18^ in hopes of extending the minibinder’s time in circulation. This fusion did not affect binding affinity or neutralization (Figure S4). With this construct, we repeated the TcsL challenge under the same conditions as before but now observed significant survival extension from a median of 4 h with the vehicle control to 8 h with the B4-M79 fusion (Figure 2A).

**Figure 2.**
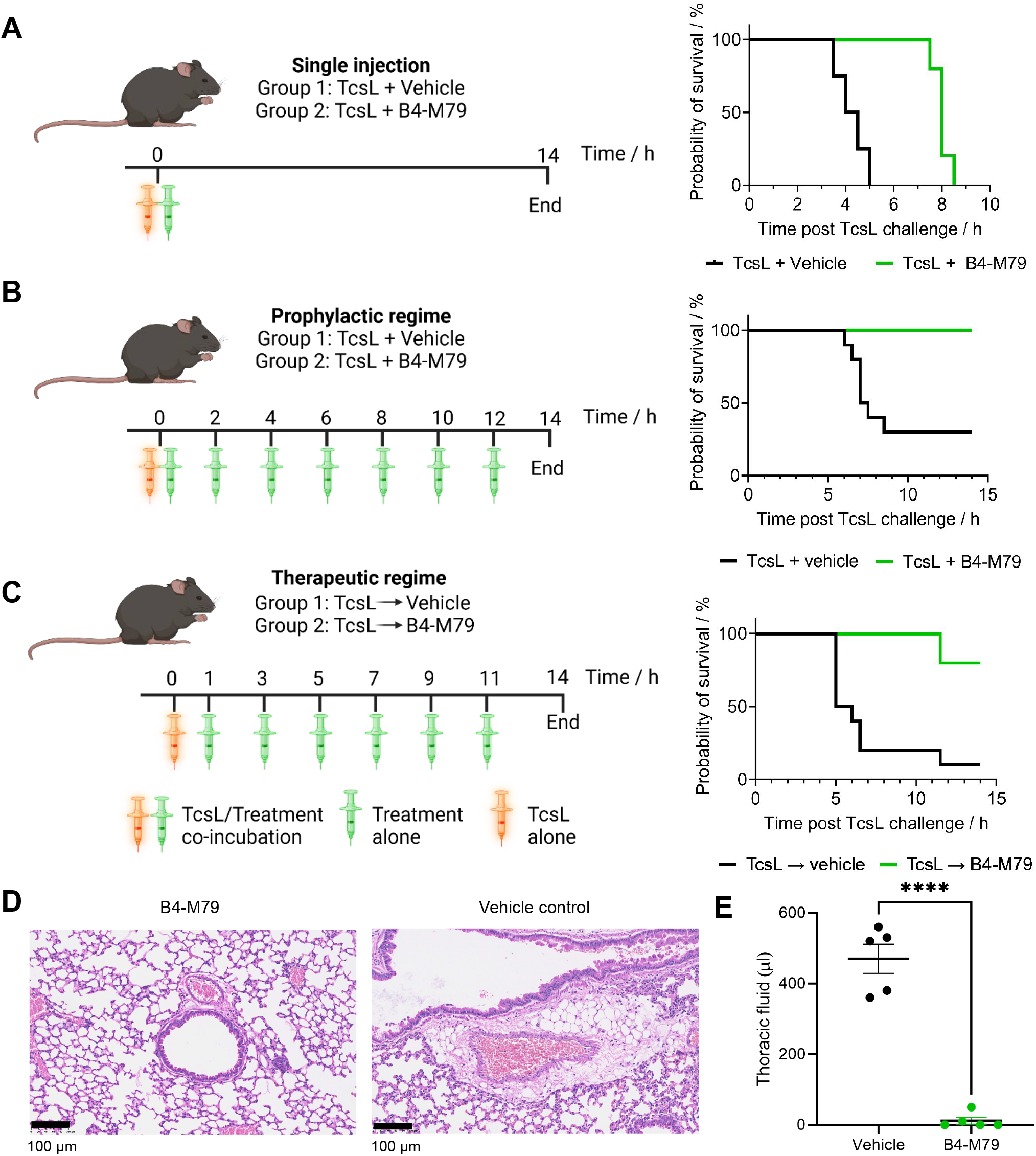
*In vivo* evaluation of inhibitors targeting TcsL in a lethal mouse challenge in three different dosing regimes. **A**. Administration of a single bolus of TcsL alone or co-incubated with a 1000x molar access of B4-M79. B4-M79 extended survival time from a median of 4.25 h to 8 h p = 0.0027 (log-rank test) **B**. Administration of TcsL alone or co-incubated with a 1000x molar excess of B4-M79, followed by dosing with the same 1000x molar excess (0.06 mg/kg) every 2 h, p = 0.0002. **C**. Administration of TcsL followed by vehicle or B4-M79 1 h later, and repeated dosing every 2 h, p = 0.0012. **D**. Example histological images of mouse lung from treatment group B with H&E stain showing greater cellular infiltrates and compressed alveolar space in the vehicle control group. **E**. Quantification of fluid build up in lung of control mice vs treated mice in group B. Lines indicate mean +/-standard error. P < 0.0001 from a two-tailed unpaired t test.

We next explored whether repeated dosing at 2 h intervals could further improve survival in both a prophylactic (co-incubation of toxin and drug prior to administration) and therapeutic (toxin administration followed by drug administration 1 h later). In the prophylactic setting, mice that received the B4-M79 therapy were completely protected from death, with little to no indication of lung edema, as assessed by the collection of fluid from the lung and histology (Figure 2B, 2D, 2E, S5). In the more stringent therapeutic setting 8/10 mice survived until the study end (14 h post toxin administration) with the other two animals exhibiting disease symptoms and becoming moribund at 11.5 h (Figure 2C, S5). In contrast, the median survival time of those receiving vehicle alone after toxin administration was 5.5 h (Figure 2C). Importantly, B4-M79 also could be lyophilized with no loss in neutralization, providing a viable pathway for long term storage (Figure S4).

TcsL is a potent cytotoxin, with lower minimum lethal doses in mice than the related toxins TcdA, TcdB, and TcsH by a factor of 10 (5 ng vs 50 - 90 ng)^19^. In humans, TcsL produced by *P. sordellii* causes a highly lethal illness with no effective treatments. The design of *de novo* inhibitors to this deadly class of toxins show that toxin-directed therapy can protect against disease by the immediate inactivation of the disease-causing molecules. In situations where patients rapidly progress towards severe illness and death, as is the case in many forms of toxic shock syndrome, antibiotics that clear the pathogen producing the toxin do not work on timescales compatible with survival. Our approach could be readily extended to other toxins implicated in toxic shock syndromes, such TSST-1 of *Staphylococcus aureus*, by blocking MHC class II and TCR binding sites on the toxic that drive non-specific immune cell activation. With only 96 designs ordered to reach a positive *in vivo* endpoint, our work highlights the democratization of protein design compared to previous efforts that required screening >100,000 designs followed by extensive experimental sequence optimization^15^, and the power of generative protein design to produce potentially life saving medicines.

## Acknowledgements

We thank Kandise VanWormer, Hernan Nunez-Ortega, Rafael Ticzon, Anusree Ravooru and Andre Dubief for lab support; Luki Goldschmidt and Patrick Vecchiato for maintaining the computational resources at the IPD; Basile Wicky and Lukas Milles for helpful discussion; and Helen Eisenach for reagent provision.

R.J.R is a Washington Research Foundation Postdoctoral Fellow. H.L. is a recipient of the Pfizer Graduate Scholarship. This work was supported by the Howard Hughes Medical Institute grant number 0001091096 (D.B.), the Department of the Defense, Defense Threat Reduction Agency grant HDTRA1-21-1-0007 (R.J.R.), The Audacious Project at the Institute for Protein Design, the Bill & Melinda Gates Foundation #INV-010680, NSERC Discovery Grant, and Canadian Institutes of Health Research Project Grant (R.A.M).

## Author Contributions

RJR, HL, RAM and DB conceptualized the project. RJR designed the binders, produced the recombinant protein, and did the binding assays with support from MG. HL did the neutralization assays. DS did the *in vivo* experiments with support from JMB. DS, RAM and DB were responsible for funding acquisition and supervision. RJR wrote the first draft of the manuscript. All authors were responsible for reviewing and editing the manuscript.

## Supplemental

### Materials and Methods

#### Design of SEMA6A-blocking minibinders

Initial backbones were generated using RFdiffusion. The target structure was from the RBD of TcsL bound to SEMA6A (6WTS^8^) (but with the receptor chain removed) and hotspot residues F1488, I1493, L1505 and I1507 with RFdiffusion. 10,000 outputs were generated and then sequence designed using ProteinMPNN and FastRelax^11,16^. One sequence from ProteinMPNN pre-FastRelax and one sequence post-FastRelax were generated for each design both at a temperature of 0.001. These designs were then filtered on the basis of predicted local distance difference test (pLDDT) > 90 and the top 48 designs with the lowest pairwise aligned error (PAE) interaction were ordered, corresponding to a cutoff of 5.7^16^.

During the second round of design, 10,000 new sequences of backbone of design F4 were generated at temperature 0.4 (no FastRelax was used here) and again filtered with 48 new designs ordered with a pLDDT cutoff of 96 and PAE interaction cutoff of 4.5.

All designed sequences for initial hits and optimization and their associated eblock gene fragment sequences are provided in the supplementary data file.

#### Expression of de novo designed miniproteins

Each design was ordered as an Integrated DNA Technologies (IDT) eblock with flanking BsaI cut sites and compatible overhangs with the LM0627 vector (Addgene 191551). Since these designs were all shorter than the 300 bp minimum, each gene was padded with arbitrary nucleotides outside of the golden gate cut sites. LM0627 encodes an MSG on the N terminus of the proteins and SNAC^20^ and 6xhis tag on the C terminus. Golden gate cloning reactions were done in 1uL total volume and all liquid handling was done by an Echo liquid handler (Labcyte). The reaction set up was:

| Reagent | Concentration | Volume (nL) |
| --- | --- | --- |
| Nuclease free water | NA | 225 |
| T4 ligase buffer | 10x | 100 |
| 300 bp eblock fragment | 4 ng/μl | 150 |
| BsaI-HFv2 | 20,000 U/mL | 50 |
| T4 ligase | 400,000 U/mL | 100 |
| LM0627 | 100 ng/μl | 325 |

Reactions were incubated at 37°C for 20 min before transformation into BL21 via 10 s heat shock at 42°C, and 1 h recovery again at 37°C in 100 μL SOC media. Then for each design, 4×1mL autoinduction media (AIM) cultures in 96-well plates were inoculated with 25 μL of each culture and incubated for 22 h at 37°C with shaking.

Cells were pelleted via centrifugation at 4000xg for 5 min and then lysed by incubation in B-PER (Thermo Fisher) with 0.1 mg/ml lysozyme, 10 μg/ml DNAse I and 1 mM PMSF for 15 min. Lysates were then cleared by centrifugation at 4000xg for 15 min and then applied to 50 μL Ni-NTA resin (Thermo Fisher) in a 96-well 25 μm plate. The resin was then washed 3x with 300 μL of wash buffer (20mM Tris, 300mM NaCl, 25mM imidazole). Protein was then eluted with 200 μL of elution buffer (20mM Tris, 300mM NaCl, 500mM imidazole pH 8). For each design, 100 μL of elute was injected onto an S75 5-150 column (Cytiva) using an AKTA autosampler (Cytiva) and underwent size exclusion chromatography in PBS (or HBS-EP+, 0.01 M HEPES pH 7.4, 0.15 M NaCl, 3 mM EDTA, 0.005% v/v Surfactant P20, in the case of samples prepared for SPR (see below)). Fractions were pooled and normalized using an OT2 (Opentrons). Overall yields were determined through the integration of the main peak from the SEC trace and aggregation state was determined based on elution volume of primary peak relative to a standard curve.

For *in vivo* experiments, expressions were done at the 50 mL scale with AIM inoculated directly from a glycerol stock and grown for 16 h at 37°C. Cells were harvested as above, resuspended in 5 mL lysis buffer (20mM Tris, 300mM NaCl, 25mM imidazole, 0.1 mg/ml lysozyme, 10 μg/ml DNAse I and 1 mM PMSF) and then sonicated at 70% amplitude, 10 s on 10 s off for 5 min. Lysates were clarified via centrifugation at 13000 x g for 15 min and supernatants applied to 0.5 mL of NiNTA resin and washed and eluted as described above (here with a 1 mL elution volume). The 1 mL elutes were injected onto a S75 10-300 Increase GL (Cytiva) for size exclusion chromatography into PBS.

#### Surface plasmon resonance

For SPR experiments, 500 response units (RU) of TcsL RBD were immobilized on a Biotin Capture chip (Cytiva) to achieve target Rmax of the analyte of 30-70 response units (RU). Designed proteins were purified into HBS-EP+ as described above. The concentrations of analyte used for each single kinetic analysis are specified in figure legends. Association occurred for 120 s at a flow rate of 30 μl/min followed by 60 s dissociation between concentrations for 6 steps of increasing concentration. Following the final injection step, dissociation was measured over 10 min - 120 min (as indicated on the SPR figure axes). Global fitting of a Langmuir 1:1 interaction model was used to determine the k_on_, k_off_, and K_D_ for each design using the Biacore 8K Evaluation software (Cytiva).

#### TcsL neutralization assays

Neutralization of TcsL was tested using HCT116 and HULEC5a cells (ATCC). Cells were seeded at a density of 4,000 cells/well in 96 well clear CellBind plates (Corning). The next day following cell attachment, minibinders were serial diluted in PBS with indicated concentration of TcsL. The solution was then added to cells and incubated for 48 hours at 37°C, 5% CO2. Cell viability was determined using alamarBlue reagent (ThermoFisher), and fluorescence was read after 2 h using a Spectramax m5 plate reader (Molecular Devices). Neutralization curves are a representative curve from 3 independent replicates while the reported IC50s are the mean from across the 3 replicates.

#### M79 fusions

B4 (sequence in supplementary data file) was produced as both an N- and C-terminal fusion to the albumin binding domain M79^18^ with 3 different linkers, GS, 2xG4S and 3xG4S. All versions of these fusions retained TcsL inhibition. The design taken forward was B4-GS-M79. The full sequence is:

MSGTEEIIKEIKKMKEKGEKASSIIDLLKKAGYSEIAKKAELAALKSENMTEAAAEKAIEELKGSQVKLEESGGGLVQAGGSLKLSCAASGSTFSSSSVGWYRQAPGQQRELVAAITSGGSTNTADSVK GRFTMSRDNAKNTVYLQMRDLKPEDTAVYYCNVAGRNWVPISRYSPGPYWGQGTQVTVSSGSGSHHWGSTHHHHHH

MSG- and -GSGSHHWGSTHHHHHH encoded by the vector LM0627.

#### *In vivo* evaluation of protection from TcsL toxic shock

Animal husbandry, ethical handling of mice and all animal work were carried out according to guidelines approved by Canadian Council on Animal Care and under protocols approved by the Centre for Phenogenomics Animal Care Committee (28-0431H). Female C67/Bl6J mice, 8 weeks of age, were intraperitoneally injected with TcsL with or without B4 (anti-TcsL minibinder) fused to M79 (albumin binding domain). Treatment-naive, littermate female mice were group-housed and randomly assigned to experimental groups. In brief, 15 ng of TcsL was mixed with a 1000-fold molar excess of B4-M97 at 4°C on a rotator for 1 h prior to injection. After intoxication, animals were monitored closely and moribund animals were euthanized. For experiments that included follow-up dosing at two hour intervals, each subsequent dose was the same as the initial dose of B4-M79, which corresponded to 0.06 mg/kg. All p values determined by Logrank test for survival curves.

For histology, lungs were harvested and fixed in 4% paraformaldehyde for 24 hours and then transferred to 70% ethanol for storage. Tissues were subsequently processed and embedded in paraffin following standard procedures at the CFIBCR Histology/Microscopy Core, University Health Network, Toronto, Canada. Paraffin blocks were sectioned at 4 μm then deparaffinized and stained for H&E using a Leica Auto Stainer XL.

---

**Figure S1.**
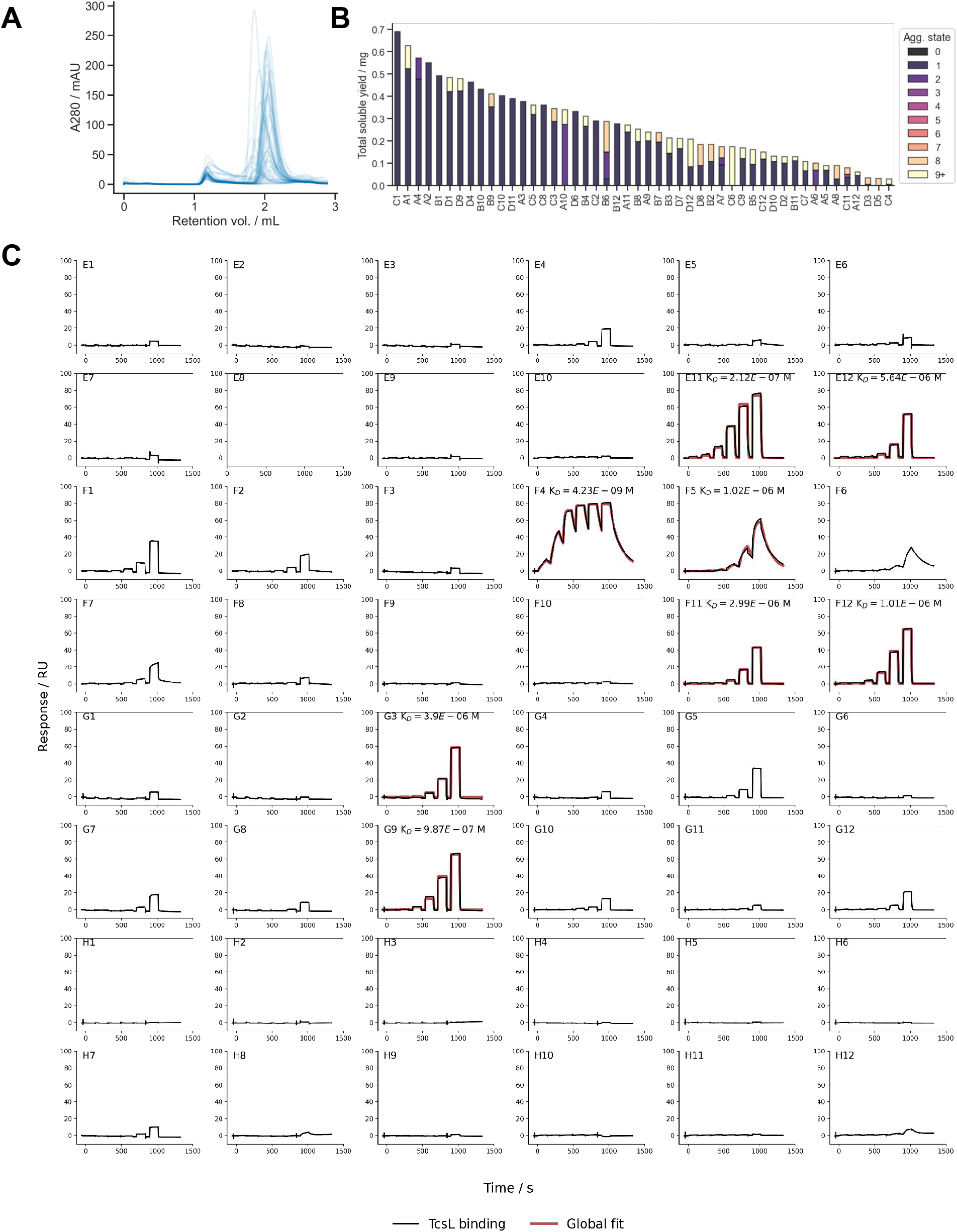
Characterization of SEMA6A-blocking miniproteins. **A**. SEC traces of all 48 designs from 4 mL cultures. **B**. Aggregation state(s) of each design based on SEC profile and molecular weight standard curve for the column. **C**. Affinity determination through SPR with the RBD of TcsL captured on the chip and a 6-step 5-fold dilution series of each miniprotein starting at 5000 nM. Global fit is shown in red while the measured data is shown in black.

**Figure S2.**
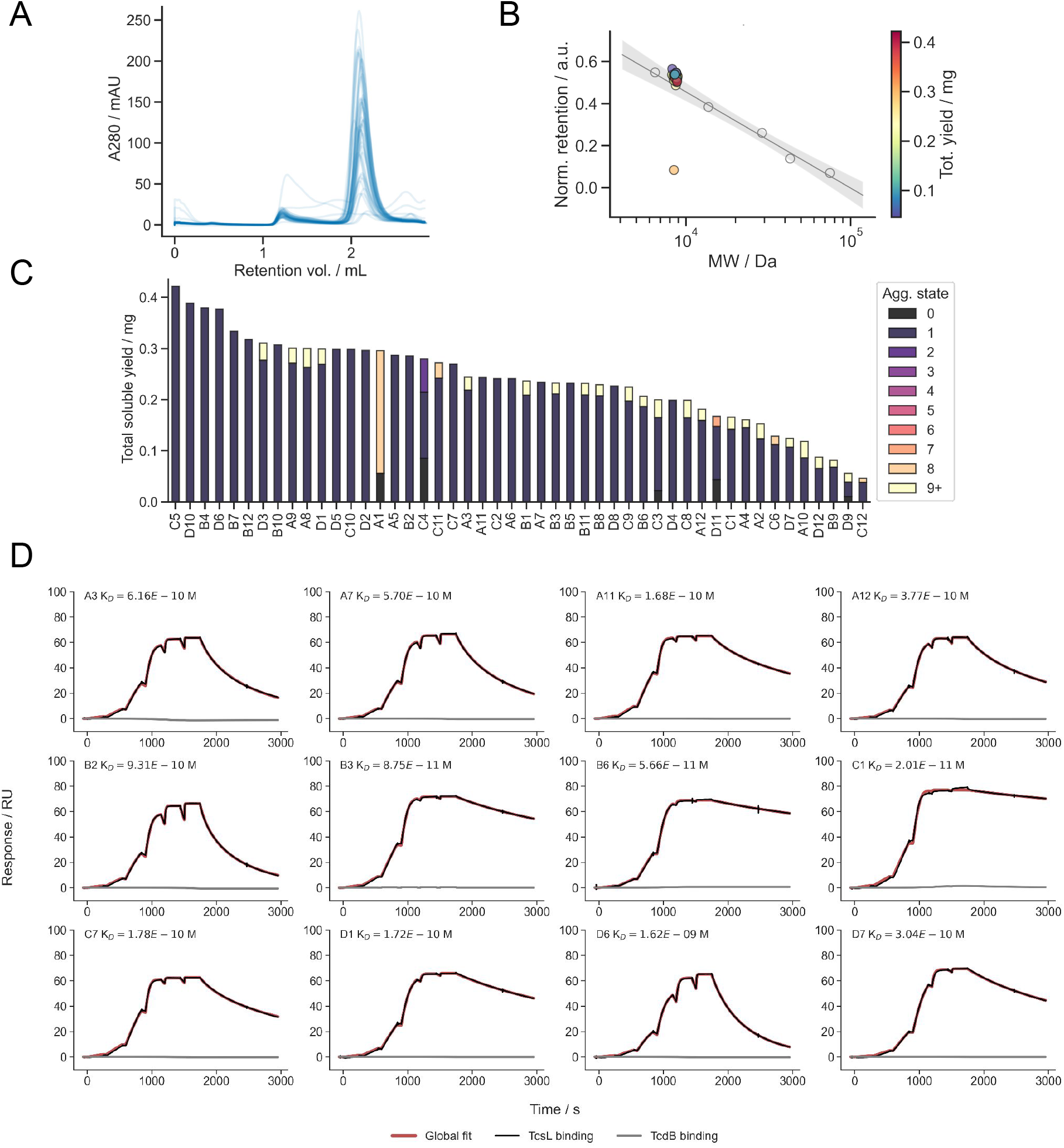
Optimization of SEMA6A-blocking miniproteins. **A**. SEC traces of all 48 designs from 4 mL culture. **B**. Molecular weight standard curve for the column to determine aggregation state from retention volume. **C**. Aggregation state(s) of each design based on SEC profile and molecular weight standard curve for the column. **D**. Affinity determination through SPR with the RBD of TcsL captured on the chip and a 6-step 5-fold dilution series of each miniprotein starting at 100 nM. Global fit is shown in red while the measured data is shown in black and TcdB binding in gray.

**Figure S3.**
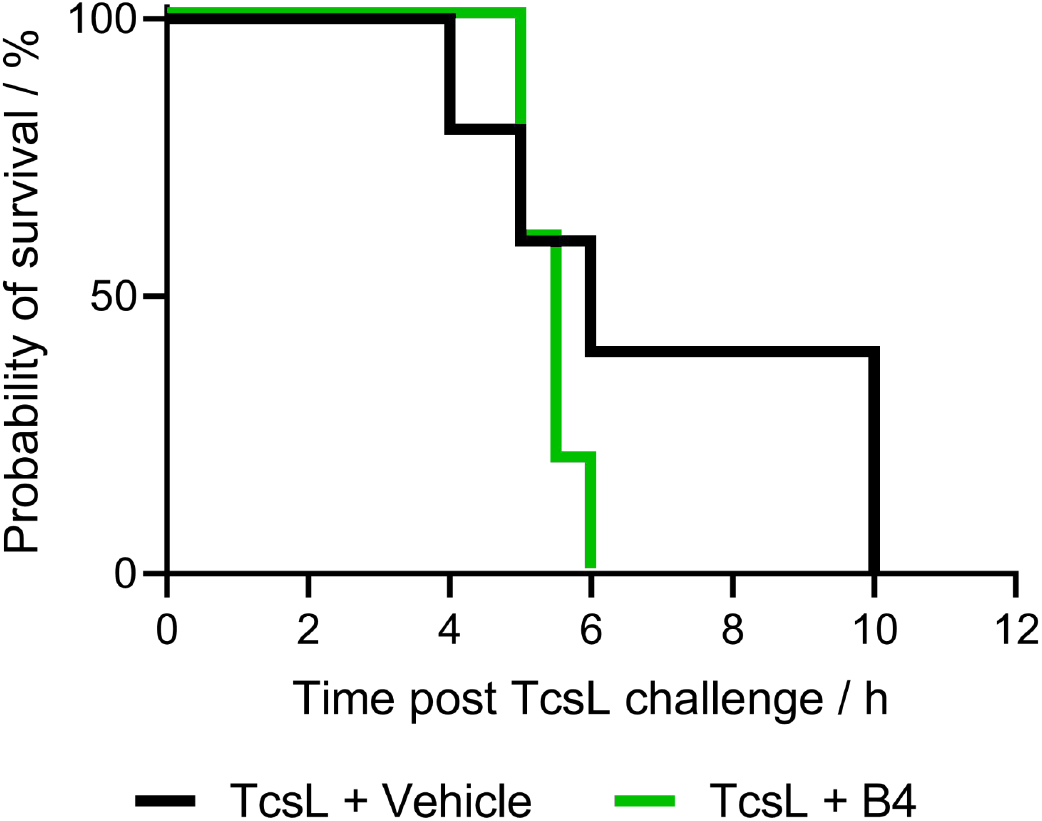
Unfused B4 monomers failed to protect against lethal toxic shock syndrome. Administration of a single bolus of 15 ng of TcsL alone or co-incubated with a 1000x molar access of B4. N = 5 in each group. No significant difference was observed between the treated and untreated groups, p = 0.27 by log-rank test.

**Figure S4.**
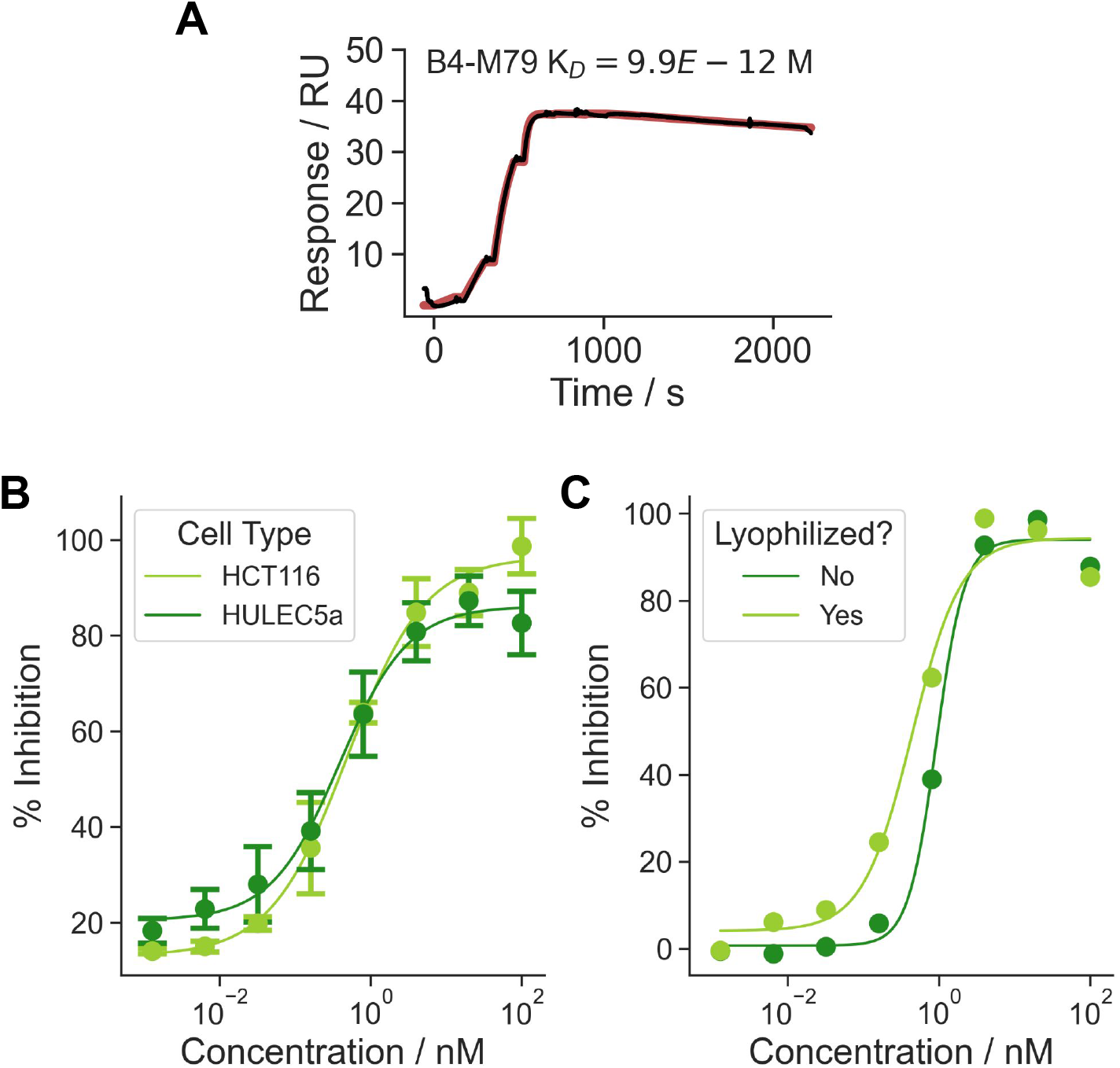
B4-M79 fusions retain binding affinity and neutralization. **A**. Single cycle kinetic analysis by SPR of B4-M79 fusion with an upper concentration of 200 nM and 6-step 5-fold dilution series. **B**. Neutralization of TcsL in HCT116 and HULEC5a cell lines in the presence of 5 µM human serum albumin with 50 pM and 0.5 pM of toxin, respectively. IC50 of 367 pM (95% CI 177 - 776 pM) for HCT116 400 pM (95% CI 140 - 963 pM) for HULEC5a. Point indicates the mean and error bars are the SEM across 3 independent replicates. **C**. Neutralization of 50 pM TcsL using HCT116 cells with lyophilized and un-lyophilized B4-M79.

**Figure S5.**
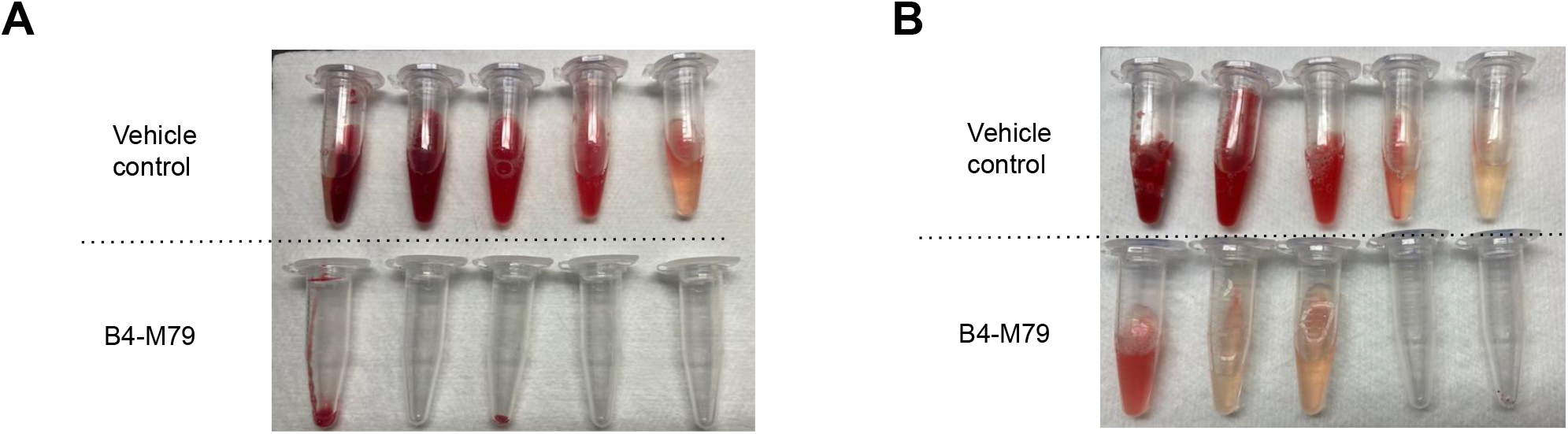
Lung edema in 5 mice treated with vehicle of B4-M79 in the prophylactic (A) or therapeutic (B) regimes.

## References

1. Fischer Marc et al. Fatal Toxic Shock Syndrome Associated with Clostridium sordellii after Medical Abortion. N. Engl. J. Med. 353, 2352–2360.

2. Aldape, M. J., Bryant, A. E. & Stevens, D. L. Clostridium sordellii infection: epidemiology, clinical findings, and current perspectives on diagnosis and treatment. Clin. Infect. Dis. 43, 1436–1446 (2006).

3. Aronoff, D. M. & Marrazzo, J. M. Infections caused by Clostridium perfringens and Paeniclostridium sordellii after unsafe abortion. Lancet Infect. Dis. 23, e48–e55 (2023).

4. Aronoff, D. M. & Ballard, J. D. Clostridium sordellii toxic shock syndrome. Lancet Infect. Dis. 9, 725–726 (2009).

5. Elkbuli, A. et al. Survival from Clostridium toxic shock syndrome: Case report and review of the literature. Int. J. Surg. Case Rep. 50, 64–67 (2018).

6. Guzzetta, M., Williamson, A. & Duong, S. Clostridium Sordellii as an Uncommon Cause of Fatal Toxic Shock Syndrome in a Postpartum 33-Year-Old Asian Woman, and the Need for Antepartum Screening for This Clostridia Species in the General Female Population. Lab. Med. 47, 251–254 (2016).

7. Hao, Y. et al. Lethal toxin is a critical determinant of rapid mortality in rodent models of Clostridium sordellii endometritis. Anaerobe 16, 155–160 (2010).

8. Lee, H. et al. Recognition of Semaphorin Proteins by P. sordellii Lethal Toxin Reveals Principles of Receptor Specificity in Clostridial Toxins. Cell 182, 345–356.e16 (2020).

9. Tian, S. et al. Genome-Wide CRISPR Screen Identifies Semaphorin 6A and 6B as Receptors for Paeniclostridium sordellii Toxin TcsL. Cell Host Microbe 27, 782–792.e7 (2020).

10. Watson, J. L. et al. De novo design of protein structure and function with RFdiffusion. Nature 620, 1089–1100 (2023).

11. Dauparas, J. et al. Robust deep learning-based protein sequence design using ProteinMPNN. Science 378, 49–56 (2022).

12. Rocklin, G. J. et al. Global analysis of protein folding using massively parallel design, synthesis, and testing. Science 357, 168–175 (2017).

13. Cao, L. et al. Design of protein-binding proteins from the target structure alone. Nature 605, 551–560 (2022).

14. Roy, A. et al. De novo design of highly selective miniprotein inhibitors of integrins αvβ6 and αvβ8. Nat. Commun. 14, 5660 (2023).

15. Cao, L. et al. De novo design of picomolar SARS-CoV-2 miniprotein inhibitors. Science 370, 426–431 (2020).

16. Bennett, N. R. et al. Improving de novo protein binder design with deep learning. Nat. Commun. 14, 2625 (2023).

17. Geny, B. et al. Clostridium sordellii lethal toxin kills mice by inducing a major increase in lung vascular permeability. Am. J. Pathol. 170, 1003–1017 (2007).

18. van Faassen, H. et al. Serum albumin-binding VH Hs with variable pH sensitivities enable tailored half-life extension of biologics. FASEB J. 34, 8155–8171 (2020).

19. Martinez, R. D. & Wilkins, T. D. Comparison of Clostridium sordellii toxins HT and LT with toxins A and B of C. difficile. J. Med. Microbiol. 36, 30–36 (1992).

20. Dang, B. et al. SNAC-tag for sequence-specific chemical protein cleavage. Nat. Methods 16, 319–322 (2019).

